# T cells are present in non-diabetic islets and accumulate during aging

**DOI:** 10.1101/2020.07.21.212977

**Authors:** Heather C. Denroche, Stéphanie Miard, Sandrine Sallé-Lefort, Frédéric Picard, C. Bruce Verchere

**Affiliations:** Canucks for Kids Fund Childhood Diabetes Laboratories, BC Children’s Hospital Research Institute, Departments of Surgery, University of British Columbia, Vancouver, British Columbia, Canada; Pathology and Laboratory Medicine, University of British Columbia, Vancouver, British Columbia, Canada; Centre for Molecular Medicine and Therapeutics, University of British Columbia, Vancouver, British Columbia, Canada; Institut universitaire de cardiologie et de pneumologie de Québec; Faculté de pharmacie, Université Laval, Québec, Québec, Canada

## Abstract

**Background:** The resident immune population of pancreatic islets has roles in islet development, beta cell physiology, and the pathology of diabetes. These roles have largely been attributed to islet macrophages, comprising 90% of islet immune cells (in the absence of islet autoimmunity), and, in the case of type 1 diabetes, to infiltrating autoreactive T cells. In adipose, tissue-resident and recruited T and B cells have been implicated in the development of insulin resistance during diet-induced obesity and aging, but whether this is paralleled in the pancreatic islets is not known. Here, we investigated the non-macrophage component of resident islet immune cells in islets isolated from C57BL/6J male mice during aging (3 to 24 months of age) and following diet-induced obesity (12 weeks 60% high fat diet). Immune cells were also examined by flow cytometry in cadaveric non-diabetic human islets.

**Results:** Immune cells comprised 2.7 ± 1.3% of total islet cells in non-diabetic mouse islets, and 2.3 ± 1.7% of total islet cells in non-diabetic human islets. In 3-month old mice on standard diet, B and T cells each comprised approximately 2-4% of the total islet immune cell compartment, and approximately 0.1% of total islet cells. A similar amount of T cells were present in non-diabetic human islets. Islet T cells were comprised of CD8-positive, CD4-positive, and regulatory T cells. Interestingly, while islet B cells and macrophage numbers were unaltered by age, the number of islet T cells increased linearly (R^2^=0.9902) with age from 0.10 ± 0.05% (3 months) to 0.38 ±0.11% (24 months) of islet cells. This increase was uncoupled from body weight, and was not phenocopied by a degree similar weight gain induced by high fat diet in mice.

**Conclusions:** This study reveals that T cells are a part of the normal islet immune population in mouse and human islets, and that they accumulate in islets during aging in a body weight-independent manner. Though comprising only a small subset of the immune cells within islets, islet T cells may play a role in the physiology of islet aging.

## BACKGROUND

Infiltration of pancreatic islets by immune cells, namely T cells, is a well known pathology of type 1 diabetes. More recently, islet inflammation has been recognized as a hallmark of type 2 diabetes (1–3), and is particularly induced by the presence of islet amyloid (4–9). The interactions between immune cells and beta cells are not solely deleterious. Healthy, non-diabetic islets contain resident macrophages, making up ≥ 90% of islet immune cells, and are present in quantities of 2 to 13 macrophages per islet (10–14). Islet macrophages play important roles in tissue homeostasis, islet development, beta cell regeneration, and physiological insulin secretion in mice (15–19). Little is known, however, about the other cells comprising resident immune cells within islets, and whether these are altered in islets under various metabolic states.

T and B cells in adipose tissue have been implicated in modulating insulin sensitivity. In obesity, B2 cells and Th1-polarized T cells accumulate in visceral adipose tissue and contribute to insulin resistance, whereas B1, B-regulatory cells, Th2 and T-regulatory cells promote insulin sensitivity, and are reduced in number and/or function in obese visceral adipose tissue (20–28). Aging is also associated with inflammation in adipose tissue promoting insulin resistance, though the changes are distinct from those of obesity. T cells accumulate in aged visceral adipose tissue whereas the total number of macrophages is relatively unperturbed (21,29–31), and unlike in diet-induced obesity, adipose tissue T regulatory cells accumulate and contribute to worsening glucose homeostasis during aging (21,29). B2 cells also accumulate in WAT with age, and contribute to insulin resistance and glucose intolerance (32). Despite the growing body of evidence that adipose-tissue T and B cells are important in glucose metabolism, the role of these cells in pancreatic islets during aging and obesity is unclear. In this study, we aimed to investigate whether lymphocytes are present in non-diabetic pancreatic islets, and whether their numbers are altered by age or obesity.

## RESULTS

### Glucose tolerance and insulin secretory capacity increase in advanced-age mice

We first examined the impact of aging from 3 to 24 months on glucose metabolism in male C57BL/6J mice. Body weight increased with age up to 12 months, but was reduced in 24-month old mice (36.1 ± 4.1 g), to a level comparable to 6-month old mice (35.4 ± 2.1 g) (Figure 1A). A decline in body weight at 2 years of age is consistent with previously published observations across many inbred mouse strains (33). Non-fasted and 6-hour fasted blood glucose levels were not altered by aging (Figure 1B,C). Aging to 6 and 12 months corresponded with moderately impaired glucose tolerance relative to 3-month old mice (Figure 1C-D), and aging to 24 months substantially lowered glucose excursions compared to all other age groups, including 3-month old mice. This finding is consistent with previous reports that, correlating with body weight, glucose tolerance is improved in advanced-aged male mice (34,35). Insulin tolerance tests indicated that insulin sensitivity cannot account for the improvement in glucose tolerance in 24-month old mice (Figure 1E). Fasted plasma insulin levels increased with age up to approximately 2.7 fold at 12 months (Figure 1F), consistent with previous observations in both mice and humans (34,36). In 24-month old mice, fasted insulin levels were similar to those of 3-month old mice. *Ex vivo* insulin secretion from isolated islets in low glucose conditions tended to increase with age up to 12 months, and glucose- and KCl-stimulated insulin secretion increased with age, with 24-month old mice displaying the highest stimulated responses compared to all other age groups (Figure 1G,H). Islet insulin content was not altered in aged mice (Figure 1I). The increased glucose- and KCl-stimulated insulin secretion in 24-month old mice suggests that stimulus-coupled insulin secretion is augmented with age, potentially accounting for improved glucose tolerance in advanced-aged mice, and supporting previous findings that aged mice have increased insulin secretion independent of insulin content (37,38).

**Figure 1.**
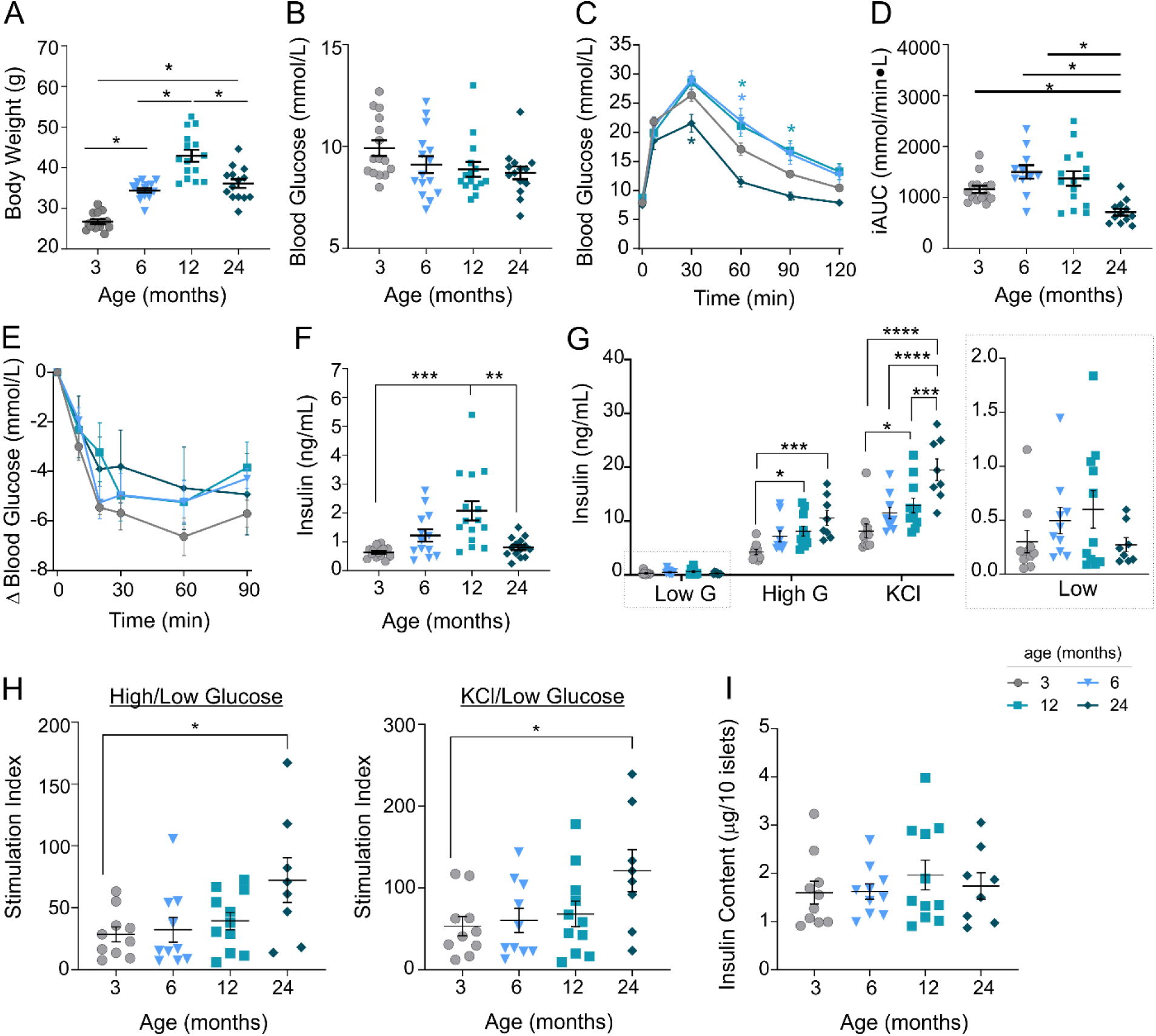
Metabolic characterization of aged mice. Metabolic measurements were performed in 3-, 6-, 12- and 24-month old C57BL/6 male mice. Non-fasted body weight (A) and blood glucose (B); glucose tolerance tests represented as glucose traces (C, n=11-15) and incremental AUC (D); insulin tolerance tests normalized to baseline (E, n=4-5), and 6-hour fasted plasma insulin (F). G-H) Insulin secretion from isolated islets, under low glucose (low G), high glucose (high G) and 30 mM KCl, represented as insulin concentration (G, with low G condition shown again on smaller scale on the right), and stimulation index relative to low glucose (H). Insulin content was measured following the final KCl stimulation (I). Data represent values for individual mice with mean ± SEM overlaid. Statistical analyses were performed via Kruskal-Wallis test with Dunn’s post-hoc test (A,D,F), or one-way ANOVA (B,G-I) or two-way repeated measures ANOVA (C,E) with Tukey’s post-hoc test (B,C,E,G-I). * p<0.05, ** *p*<0.01, *** *p*<0.001, **** *p*<0.0001 vs 3-month old unless other comparison specified.

### T cells accumulate in aging islets

We next assessed the effect of aging on islet-resident immune populations. Islets from C57BL/6J mice of the four age groups were isolated and hand-picked to purity. Islets from a minimum of five mice were pooled per sample, to obtain a sufficient number of islet immune cells, and dispersed into single-cell suspensions for analysis by flow cytometry (Figure 2A). Islet cells were gated on singlets and viability prior to analysis of immune cell populations (Figure 2B). Immune cells (CD45+) accounted for 2.7 ± 1.3% of viable islet cells in 3-month old mice to 3.5 ± 0.9% in 24-month old mice (Fig. 2C). As we were initially interested in resident islet B-cell populations, islet cells were additionally stained for CD19, along with CD23 and CD5 to differentiate B-cell subsets. Approximately 90% of islet immune cells were negative for CD19 and CD5 (Figure 2B), consistent with reports that the vast majority of resident islet immune cells are macrophages (10,11,39). CD19+ cells were negative for CD5, consistent with B2 cells (Figure 2B), and present at a frequency of approximately 0.1% of islet cells, which was not altered by age (Figure 2D).

**Figure 2.**
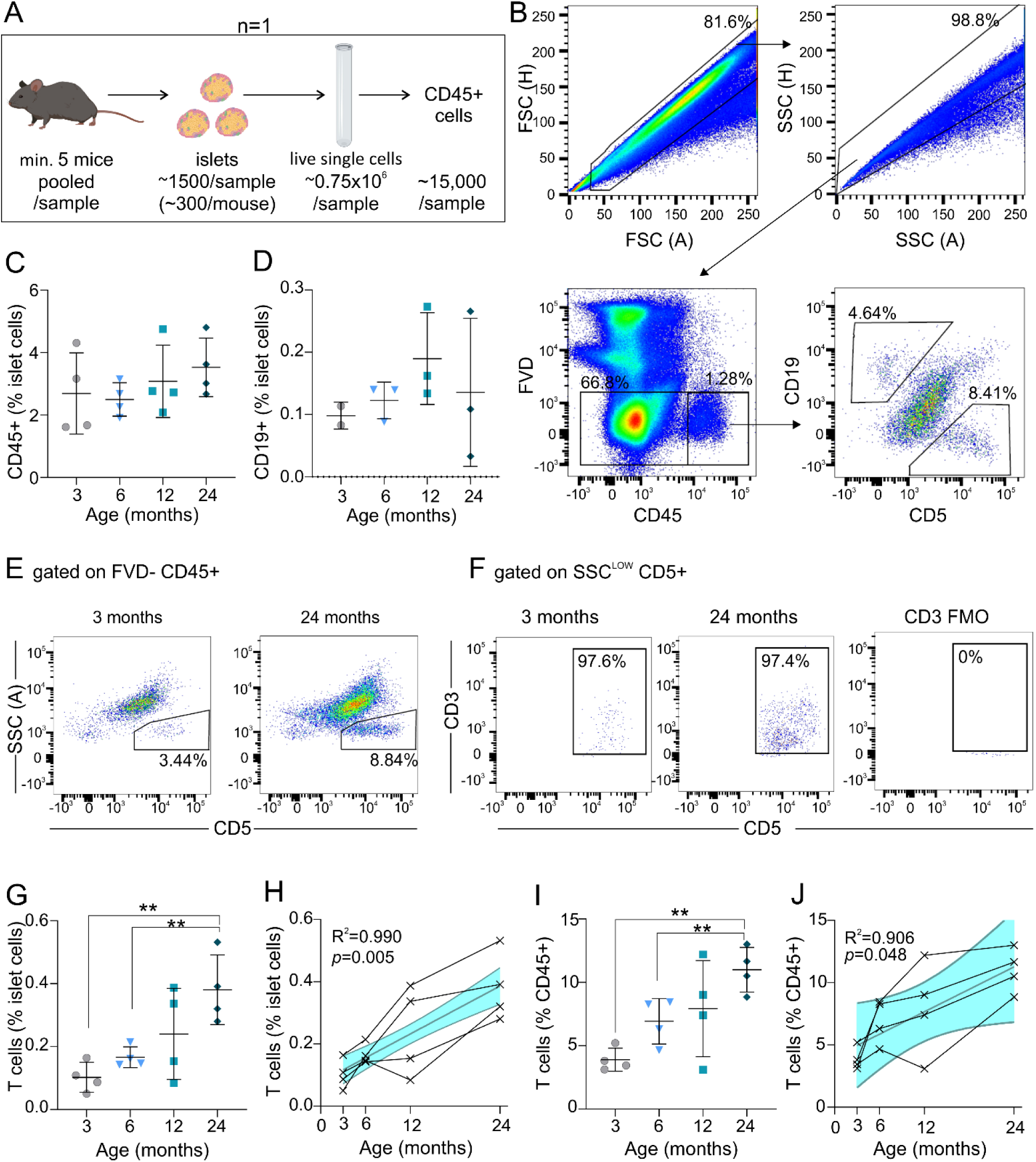
T cells accumulate in mouse islets during aging. Islets were isolated from C57Bl/6J male mice aged 3, 6, 12 and 24 months for analysis by flow cytometry. A) Schematic of flow cytometry protocol; islets from a minimum of 5 mice were pooled per sample to obtain sufficient dispersed islet CD45+ cells for analysis. Each pooled sample of ≥5 mice is considered one individual biological sample (n=1). B) Cells were gated on forward-scatter (FSC), side-scatter (SSC), viability (FVD-), CD45+, and subsequently CD19 or CD5. Data are from a representative 6-month old mouse islet sample. C-D) CD45+ and CD19+ cells expressed as a percent of total viable islet cells. E) Representative plots of SSC^low^CD5+ populations from 3-month and 24-month old mouse islets. F) Cells gated on SSC^low^CD5+ are positive for CD3+, box represents CD3+CD5+ cells. G-J) T cells expressed as a percent of total viable islet cells (G-H) or CD45+ islet cells (I-J). Linear regression (H,J) analysis shows 95% confidence interval in blue and Pearson correlation coefficient, connecting lines between data points are from a given cohort of mice run in one independent experiment. Data represent independent biological samples, each run as an independent experiment, with mean ± SD overlaid (C,D,G-J), and were analyzed by one way ANOVA with Dunnett’s T3 multiple comparisons test (C,G,I) or Kruskal-Wallis test with Dunn’s multiple comparisons test (D). * p<0.05, ** *p*<0.01, *** *p*<0.001, **** *p*<0.0001. Numbers in FACS plots represent the percent of cells in each selection as a function of the parent population.

Surprisingly, in addition to B cells, we observed that islets contained a distinct population of CD19-CD5+ cells (Figure 2B). CD19-CD5+ cells comprised 3.9 ± 0.9% of islet immune cells in 3-month old mice, had a side scatter profile (SSC^low^) consistent with lymphocytes, and accumulated with age (Figure 2E-G). As CD5 expression is limited to B1a cells and T cells, we suspected that these were islet-resident T cells. Indeed, CD19-CD5+ cells expressed the T cell co-receptor, CD3, in all age groups (Figure 2F). Islet T cells showed a positive linear correlation with age, comprising 0.10 ± 0.05% of islet cells at 3 months and rising approximately 4-fold to 0.38 ±0.11% of islet cells at 24 months of age (R^2^=0.9902, *p*=0.0049) (Figure 2G-H). This increase was due to a specific accumulation in T cells, as the frequency of T cells relative to islet immune cells also increased with age from 3.9 ± 0.9% of CD45 cells to 11.0 ±1.8% of CD45+ cells (R^2^=0.9059, *p*=0.048) (Figure 2I-J).

We next assessed the characteristics of islet T cells by FACS sorting islet cells from an additional cohort of 3- and 24-month old mice. Single, viable islet cells were gated on CD45 and subsequently on CD19 and CD3 staining (Figure 3A). Again, CD3+ cells were significantly increased in the islets of 24-month old mice compared to 3-month old mice (0.61 ± 0.08% vs 0.08 ± 0.03% of islet cells) (Figure 3A-B) whereas there were no differences in islet B cells or CD3-CD19-cells in islets with age (Figure 3B). There was no increase in CD3+ cells in the spleens of aged mice, consistent with previous reports (30) (Supplemental Figure 1A). Cell profile analysis (Nsolver4.0) of sorted islet CD3+ cells confirmed a statistically significant T cell transcript profile (Supplemental Figure 1B). Furthermore, islet CD3+ cells expressed multiple T cell specific transcripts, including Cd3d, Cd3e, Cd3z, Cd4 and Cd8, none of which were significantly altered by age (Figure 3C), suggesting that similar islet T cell subsets persist across age groups. We subsequently examined islet T cell subsets from approximately 1 year-old mice by flow cytometry (Figure 3D). CD19-CD3-cells comprised the majority of islet immune cells, and were positive for CD11b, consistent with islet macrophages. Islet CD3+ cells contained a mixture of CD8+, CD4+ and regulatory T cell subsets: 29.3 ± 0.3% were CD8+ and 38 ± 3% were CD4+, and 18.4 ± 0.3% of CD4+ cells were FoxP3+, comprising 7.1 ± 0.6% of the islet T cell population. An additional 32 ± 2% of T cells were double negative (CD8-CD4-). This was distinct from splenocyte T cell subsets, which were comprised of 45.9% CD8+ cells and 42.7% CD4+ cells, 10.8% of which were FoxP3+ (Supplemental Figure 1C).

**Figure 3.**
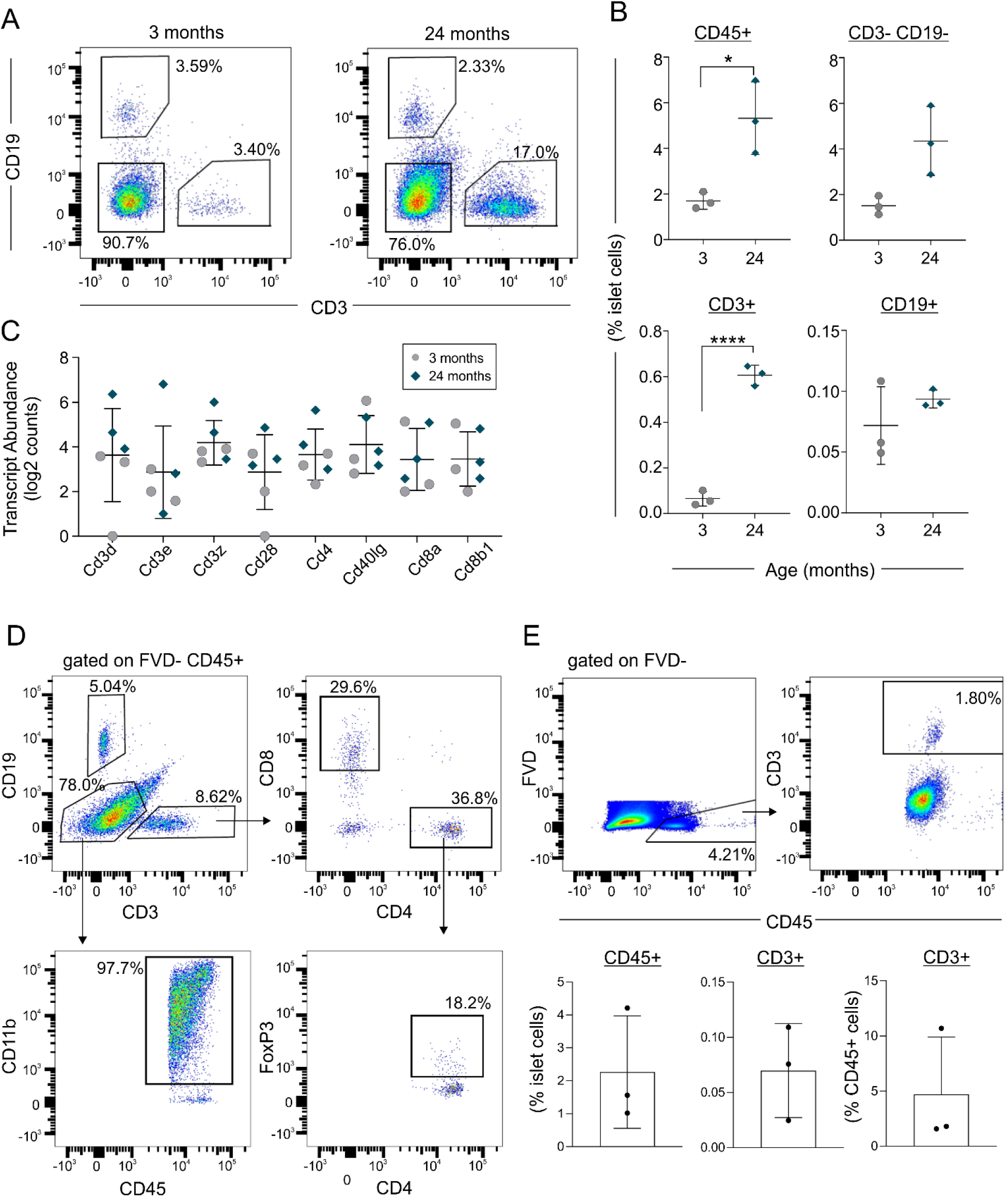
Characterization of islet T cells in non-diabetic mice and humans. A-C) Islets were isolated from 3- and 24-month old mice. Data represent 3 samples per age group, with islets from 5-7 mice pooled per sample. A) Islet cells gated on FSC, SSC, viability (7AAD-) and CD45+, representative of 3 independent samples. B) Quantification of CD45+ cells, and immune cell subsets in islets from 3- and 24-month old mice, data represent mean ± SD, and were analyzed by unpaired T-test. C) Transcript abundance of sorted CD3+ islet cells, in 3-month samples (black) and 24-month samples (blue), data represent mean ± SD. D) Islets were isolated from 10-month old mice (10 mice pooled per sample) for analysis of islet T cell subsets by flow cytometry. Cells were gated on FSC, SSC, viability (FVD-) and CD45+, and subsequently CD19, CD3, CD11b, CD8, CD4, and FoxP3; data are representative of 2 samples, islets from 10 mice per sample. E) Islets from 3 non-diabetic, cadaveric human donors were dispersed for analysis. Cells were gated on FSC, SSC and viability (FVD-) (Supplemental Figure 2). Representative data showing CD45+ cells as a function of viable islet cells (top left), CD3+ cells as a function of CD45+ cells, and quantification of islet immune populations from the 3 samples (bottom), shown as mean ± SD. Numbers in FACS plots represent the percent of cells in each selection as a function of the parent population. * p<0.05, *\*\* p*<0.01, *** *p*<0.001, **** *p*<0.0001.

### T cells are present in non-diabetic human islets

To determine whether the presence of T cells in non-diabetic islets was generalizable to humans, we performed flow cytometric analysis on human islets from 3 cadaveric, non-diabetic human donors (Supplemental Table 2). Similar to mice, all human islet preparations contained a small proportion of CD45+ cells, comprising approximately 2.3 ± 1.7 % of islet cells (Figure 3E). Furthermore, all samples contained a population of CD3+ cells within pancreatic islets, at a frequency of 0.070 ± 0.043% of total live islet cells (Figure 3E), comparable to frequencies we observed in young adult mice.

### Islet T cells are not increased by diet-induced obesity

To determine whether the accumulation of intra-islet T cells was specific to aging, or associated generally with obesity or increasing body weight, we next examined islet T cells in diet-induced obese mice. Mice were fed a high fat diet (HFD) for 12 weeks (up to 4 months of age) and compared to age-matched, low fat diet (LFD)-fed mice. This duration of HFD increased body weight by 25% relative to LFD-fed controls (39 ± 3 vs 31 ± 2 g) (Figure 4A), a comparable increase to the ∼28% increase observed in 6-month relative to 3-month old mice (34 ± 2 g vs 27 ± 2 g) (Figure 1A). In terms of absolute body weight, HFD-fed mice were in the range of 12-month (43 ± 6 g) and 24-month (36 ± 4 g) old aged-mice (Figure 1A). Non-fasted and fasted blood glucose were not altered by HFD relative to LFD (Figure 4B-C), though glucose tolerance was robustly impaired, as expected (Figure 4C-D). HFD-fed mice were also insulin resistant compared to LFD-fed controls (Figure 4E-G), and had a ∼2.6 fold increase in plasma insulin levels (Figure 4H), similar to the ∼2.7 fold increase in insulin in 12-month relative to 3-month old mice (Figure 1F). Thus, HFD achieved similar increases in body weight and insulin levels to 12 months of aging but resulted in a more dramatic impairment in glucose tolerance. T and B cells were present in islets at frequencies of ∼1% of immune cells in LFD-control mice, comparable to frequencies in 3-month old mice (Figure 4I-J). Despite comparable increases in body weight and insulin to 12 months of aging, HFD-fed mice showed no increase in islet CD3+ cells. The frequency of CD45+ cells in islets, as well as the number of CD19-CD3-cells was also unaffected by 12 weeks of high fat diet (Figure 4J). Collectively, these data show that T cells are present in islets in the absence of autoimmune diabetes, and that there is a distinct increase in islet T cells during aging that cannot be accounted for by obesity.

**Figure 4.**
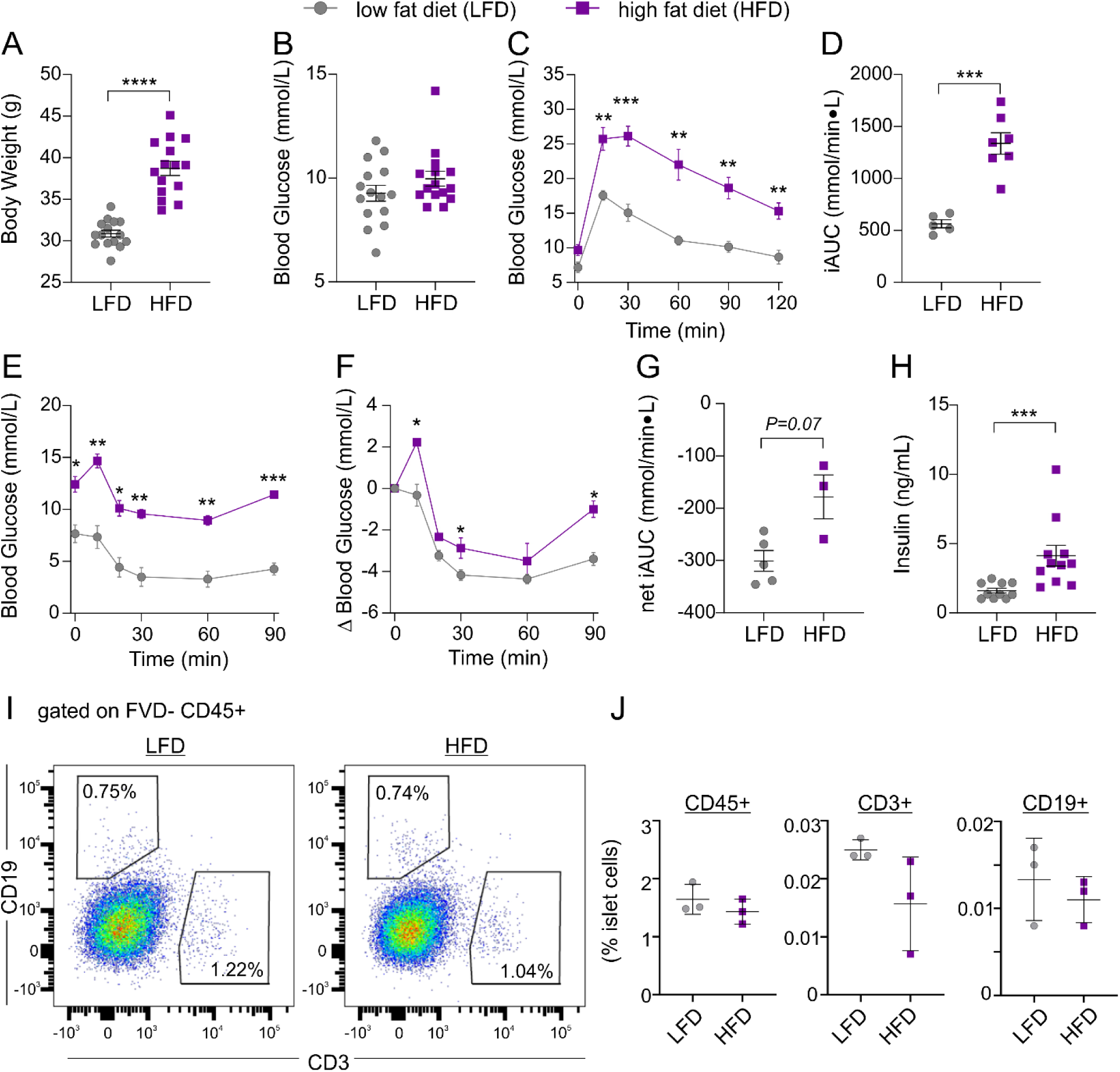
Diet-induced obesity does not cause T cell accumulation in islets. C57BL/6J mice were fed a high fat diet (HFD, purple squares) or low fat diet (LFD, grey circles) for 9-12 weeks from 6 weeks of age. Non-fasted body weight (A) and blood glucose (B) after 9 weeks of diet. Glucose tolerance tests (C-D, n=5-7), shown as glucose traces (C) and incremental AUC (D), and insulin tolerance tests (E-G, n=3-5) at 10 and 11 weeks of diet, shown as raw blood glucose (E), delta blood glucose (F), and net AUC relative to baseline (G). H) Fasted plasma insulin at 11-12 weeks of diet. Data represent mean ± SEM, and were analyzed by Student’s T-test with Welch’s correction (A,D,G), Mann-Whitney test (B,H), or two-way repeated measures ANOVA with Sidak’s multiple comparisons test (C,E,F). I-J) Islets were isolated from LFD- and HFD-fed mice after 12 weeks of diet (n=3 samples, 5 mice pooled per sample), dispersed, and gated on FSC, SSC, viability (7AAD-) and CD45+, and subsequently CD19 and CD3. I) Representative data for LFD- and HFD-mouse islets. J) Quantification of islet immune cell populations as a function of viable islet cells, data represent mean ± SD and were analyzed by Mann-Whitney test. Numbers in FACS plots represent the percent of cells in each selection as a function of the parent population.

## DISCUSSION

In this study we found that T cells contribute to the normal islet immune cell repertoire in non-diabetic mice and humans. While macrophages comprise the majority of islet immune cells (10–14,40), B and T cells each comprise up to approximately 5% of islet immune cells in young adult mouse islets. Islet T cells were comprised of a mixture of CD4+, CD8+ and FoxP3+ subsets. We also found that islet T cells accumulated within islets during aging in mice. Islet T cells increased by 4- to 8-fold from 3 to 24 months of age, and this increase was specific to T cells, with no changes in islet macrophages or B cells. Moreover, an age-related accumulation of T cells was not observed in spleen, suggesting this phenomenon is islet-specific. In addition, the particular frequencies of T cell subsets present within islets appeared distinct from those in the spleen. Finally, islet T cells were not increased by similar increases in body weight brought about by diet-induced obesity in young mice, suggesting that islet T cell accumulation is a distinct process of aging. Thus, we postulate that T cells play a role in islet biology (in the absence of autoimmune diabetes), and particularly, in adaptive changes to the pancreatic islet during aging, though a functional role for islet T cells was not tested in this study.

Our findings contrast some studies that have claimed a lack of T cells in non-diabetic mouse islets. Though scarce relative to islet macrophages, our study clearly shows a small, yet persistent population of T cells in both mouse and human islets; this is consistent with a small number of studies that have reported T cells in non-diabetic islets (2,41–43), but currently the contribution of this population to resident islet immune cells is not widely recognized. At approximately 0.1% of islet cells in young adult mice, there would be approximately one T cell present in an islet comprised of 1000 cells, which would increase to approximately 4 to 8 T cells in a similar sized islet in 2-year old mice. This does not simply reflect the expected increase in islet size or the number of total islet cells with age, as the increase is observed as a percent of total islet cells. Despite the paucity of T cells within pancreatic islets, these cells may still produce sufficient levels of local cytokines within the islet to influence the islet environment. Such is the case for islet macrophages, comprising only 2-10 cells per islet, which are the major source of islet IL-1β (6,15), and play key roles in islet development and beta-cell adaptation and expansion (15–19,40,44). Islet-resident group 2 innate lymphoid cells, another rare immune population in islets, have also been shown to influence insulin secretion, indirectly via islet macrophages (42). These examples underline how a small number of cells can have a substantial impact on islet physiology.

The increase in T cells – but not other immune cells – within aging islets, suggests these cells may play a specific aging-related role in pancreatic islets. Furthermore, the aging-induced accumulation of islet T cells is not recapitulated by a similar degree of weight gain in diet-induced obese mice, and thus is not driven by obesity. This parallels the divergent immune cell profiles in adipose tissue of aged versus obese mice (31). Similarly, the metabolic outcomes of aging and diet-induced obesity in mice are different, with aging resulting in increased insulin secretion and ultimately improved glucose tolerance in advanced age mice, whereas diet-induced obesity impairs insulin secretion and glucose tolerance. Thus, the immunologic and metabolic state of aging is distinct from that of obesity, and it is plausible that, like in adipose tissue, T cell accumulation in islets contributes to the regulation of glucose homeostasis during aging. The function of islet T cells during aging, was not examined in this study due to the large number of mice that would be required, but warrants further investigation. Indeed, the correlation between the progressive increase in insulin secretion during aging, observed in this study and others (34,35,37,38,45), and the progressive accumulation of islet T cells reported here, is intriguing. Interestingly, in pre-diabetic NOD mice, the autoreactive T cells that home to islets have been shown to promote beta cell proliferation through production of cytokines IL-2, IL-6, IL-10, CCL3, and CCL5 (46). The rise in islet T cells may also be indicative of an immune response to senescent beta cells, which accumulate in insulin resistant states including aging and obesity (47).

## CONCLUSIONS

Collectively, this study demonstrates that T cells are part of the normal immune population of pancreatic islets in non-diabetic mice and humans, and that their numbers accumulate during aging in mice. The function and role of this unique islet immune population, particularly during aging, is an interesting avenue of future study.

## METHODS

### Animals

Male C57BL/6J mice were obtained from Jackson Laboratories (Strain #000664, Jackson Laboratories, Bar Harbor, ME) and aged up to 24 months at the Institut universitaire de cardiologie et de pneumologie de Québec-Université Laval (Québec, Canada) as part of the aging mice colony of the Québec Network for Research on Aging (RQRV). Mice were housed under 12-hour light 12-hour dark conditions at room temperature, with *ad libitum* access to regular chow (NIH-31 Teklad #7917, Envigo) and water. Mice were transported at 3, 6, 12 or 24 months to BC Children’s Hospital Research Institute (Vancouver, Canada) for all experiments, where mice were housed under 12-hour light 12-hour dark conditions at room temperature, with *ad libitum* access to chow (Teklad #2918, Envigo) and water. Mice were acclimated for one week following arrival prior to any experimentation. High fat diet fed mice (Jackson Laboratories, cat#380050) were fed a 60% fat diet (#D12492, Research Diets, New Brunswick, USA) from 6 weeks of age. Chow-fed controls (Jackson Laboratories, cat#380056) were fed 10% fat diet (#D12450B, Research Diets) from 6 weeks of age. All animal procedures were approved by the Animal Care Committees at the University of British Columbia and Université Laval and carried out in accordance with the Canadian Council on Animal Care guidelines.

### Metabolic Tests

Blood glucose was measured via tail poke and measured with a OneTouch Ultra Glucometer (Life Scan Inc, Burnaby, Canada). For glucose tolerance tests, mice were fasted for 6 hours and subsequently injected intraperitoneally with 2 g/kg (aging study) or 1.5 g/kg (diet study) body weight glucose, and blood glucose measured at indicated time points as above. Blood was collected from the saphenous vein after a 6-hour fast for the measurement of plasma insulin via Stellux Chemiluminescent Rodent Insulin ELISA (#80-INSMR-CH01, Alpco, Salem). For insulin tolerance tests, insulin (Novolin GE Toronto, Novo Nordisk) was injected at the 0.65 U/kg (aging study) or 0.35 U/kg (diet study) and blood glucose measured at indicated time points.

### Flow cytometry and cell sorting

Islets were isolated via collagenase injection into the pancreatic duct, and dispersed for flow cytometry as previously published (10). Briefly, islets were harvested from mice and hand-picked to purity, and subsequently exocrine-free islets were selected and dispersed with 0.02 % trypsin for 3 min at 37 °C. Dispersions were stopped by addition of 7 mL of FACS buffer containing 2% FBS on ice. Dispersed cells were incubated with FcR block (Thermo Fisher Scientific), 10 min prior to addition of antibodies (Supplemental Table 1) for 30 min. 7-AAD or fixable viability dye eFluor780 (Thermo Fisher Scientific) served as a live-dead stain. Flow cytometry was conducted on a LSRFortessa (BD Biosciences, San Jose, USA) or BD LSR II (BD Biosciences). Cells were sorted on a BD FACS ARIA II (BD Biosciences) into PBS containing 2 % FBS.

### Transcript analysis

FACS-sorted islet cells were washed, and lysed in a minimum of 5 µL Cells-to-CT buffer containing 1% DNAse I (Taqman Gene Expression Cells-to-CT Kit, Thermo Fisher Scientific), or a volume adjusted for a final cell concentration of ∼1000 cells/µL for 5 min at RT, and stopped by addition of 10% Stop Solution, according to manufacturer instructions. Transcriptome analysis was performed via Nanostring nCounter XT (Nanostring Technologies, Seattle, USA) with the XT_PGX_MmV1_Immunology Code Set. Due to low cell input from 3-month old islets, pre-amplification of RNA was performed via Nanostring Low RNA Input Kit following manufacturer instructions. Briefly, RNA was diluted 1:2 in water, and 4 µL sample was reverse transcribed, followed by 8 rounds of amplification with M Immunology v1 Primers. All other samples were assessed without pre-amplification. Cell lysate or pre-amplified cDNA (5 µL) was added to hybridization mixtures for 18 h at 65 ° C with a 70° C-heated lid. Gene expression was measured with a Nanostring SPRINT Profiler, and data were analyzed via Nsolver 4.0 software.

### Glucose-stimulated insulin secretion

Duplicates of 10 size-matched islets per mouse were rested in Krebs-Ringer Bicarbonate buffer (129 mmol/L NaCl, 4.8 mmol/L KCl, 1.2 mmol/L MgSO_4_, 1.2 mmol/L KH_2_PO_4_, 2.5 mmol/L CaCl_2_, 5 mmol/L NaHCO_3_, 10 mmol/L HEPES, 0.5 % bovine serum albumin) containing 2.8 mM glucose for 1 hour at 37°C, and were sequentially incubated for 45 min in KRB containing 2.8 mM (low) glucose, then 16.7 mmol/L (high) glucose, and then 30 mmol/L KCl (103.8 mmol/L NaCl, 30 mmol/L KCl, 1.2 mmol/L MgSO_4_, 1.2 mmol/L KH_2_PO_4_, 2.5 mmol/L CaCl_2_, 5 mmol/L NaHCO_3_, 10 mmol/L HEPES, 0.5 % bovine serum albumin, 2.8 mmol/L glucose). Cell supernatants were collected for measurement of insulin secretion, and islets were lysed in acid:ethanol for insulin content. Insulin was measured via Chemiluminescent Rodent Insulin ELISA (#80INSMR-CH01, Alpco).

### Human islets

Human islets from cadaveric, non-diabetic donors (donor characteristics in Supplemental Table 2) were supplied by the Alberta Diabetes Institute (Edmonton, Canada). Upon receipt, islets were hand-picked to 99% purity, and were cultured in CMRL (supplemented with 100 U/mL penicillin, 100 µg/mL streptomycin, 0.05 mg/mL Gentamicin, and 2 mmol/L glutamax) containing 11 mM glucose at 25°C overnight prior to dispersion for FACS. Procedures using human islets were approved by and performed in accordance with guidelines of the Clinical Research Ethics Board of the University of British Columbia.

### Statistical analyses

With the exception of transcriptome analyses, all statistical analyses were performed using GraphPad Prism 8.0 (GraphPad Software, La Jolla, USA). Data were tested for normality by Shapiro-Wilk test. For comparisons of only two groups, normally distributed data were analyzed by t-test with Welch’s correction whereas non-normal data were analyzed by Mann-Whitney test. For comparisons of more than two groups, normally distributed data were analyzed by one-way ANOVA and non-normal data analyzed by Kruskal-Wallis test. For repeated measures of two or more groups (as in GTTs and ITTs) data were analyzed by two-way repeated measures ANOVA with Geisser-Greenhouse correction for non-sphericity. Data in all figures and text are represented as mean ± SD unless otherwise specified.

## ABBREVIATIONS

(FSC): Forward-scatter
(GTT): Glucose tolerance test
(HFD): High fat diet
(ITT): Insulin tolerance test
(LFD): Low fat diet
(SSC): Side-scatter

## DECLARATIONS

### Availability of data and materials

Open source data on each human islet preparation can be obtained at www.isletcore.ca.

### Authors’ contributions

Studies were conceived and designed by HCD, SSL, SM, FP and CBV. HCD performed data acquisition and analysis and wrote the manuscript. SSL and SM contributed to data acquisition. SSL, SM, FP, CBV edited the manuscript and approved the final version. CBV is the guarantor of this work.

## Acknowledgements

The authors acknowledge the support of the Quebec Network of Research on Aging, which allowed access to its colony of aging mice, and the Alberta Diabetes Institute for supplying human islet preparations. The authors also wish to thank the following colleagues at BC Children’s Hospital Research Institute for their research assistance: D.L. Dai, G. Soukhatcheva, M. Komba for assistance with islet isolations, N. Kim for assistance with GTTs, and D Wu and L. Xu for assistance with antibody panels and FACS.

## SUPPLEMENTAL INFORMATION

**Supplemental Table 1.** Antibodies for flow cytometry.

| **Target** | **Clone** | **Catalogue** | **Dilution** |
| --- | --- | --- | --- |
| **Mouse islets** |  |  |  |
| **CD45** | 30-F11 | 48-0451-82 | 1:100 |
| **CD19** | eBio1D3 | 17-0193-82 | 1:100 |
| **CD5** | 53-7.3 | 11-0051-82 | 1:150 |
| **CD11b** | M1/70 | 12-0112-82 | 1:100 |
| **CD3** | 17A2 | 564009 | 1:100 |
| **CD4** | GK1.5 | 100453 | 1:100 |
| **CD3e** | 145-2C11 | 25-0031-82 | 1:25 |
| **CD8a** | 53-6.7 | 100761 | 1:100 |
| **FoxP3** | FJK-16s | 11-5773-82 | 1:25 |
| **Human islets** |  |  |  |
| **CD45** | HI30 | 48-0459-42 | 1:15 |
| **CD3** | UCHT1 | 17-0038-42 | 1:20 |

**Supplemental Table 2.** Human islet donor characteristics. This table was prepared following recommendations as per: Hart N.J., Powers A.C. 2019. Use of human islets to understand islet biology and diabetes: progress, challenges and suggestions. Diabetologia 62:212–222. Open source data on each islet preparation can be obtained at www.isletcore.ca.

| Islet preparation | 1 | 2 | 3 |
| --- | --- | --- | --- |
| **Donor demographics** |  |  |  |
| Unique identifier | R310 | R314 | R317 |
| Age (years) | 25 | 31 | 54 |
| Sex (M/F) | M | F | M |
| BMI (kg/m<sup>2</sup>) | 26.4 | 30.3 | 26.4 |
| HbA<sub>1c</sub> | 5.4 | 5.0 | 5.1 |
| Cause of death | NDD-neurological | NDD-neurological | NDD-neurological |
| Diabetes? (Y/N) | N | N | N |
| **Pancreas** |  |  |  |
| Origin/source | Alberta Diabetes<br>Institute Islet Core | Alberta Diabetes<br>Institute Islet Core | Alberta Diabetes<br>Institute Islet Core |
| Cold ischaemia time (h) | 11.25 | 14.75 | 21 |
| **Islet handling and use** |  |  |  |
| Origin/source | Alberta Diabetes<br>Institute Islet Core | Alberta Diabetes<br>Institute Islet Core | Alberta Diabetes<br>Institute Islet Core |
| Isolation centre | Alberta Diabetes<br>Institute Islet Core | Alberta Diabetes<br>Institute Islet Core | Alberta Diabetes<br>Institute Islet Core |
| Estimated purity (%) | 90 | 80 | 90 |
| Total culture time (h) | ~96 | ~48 | ~96 |
| Functional measurement<br>(Stimulation index mean<br>10 mM to 1 mM) | 14.07 | 3.62 | 11.9 |
| Experimental islet use<br>(including in which<br>experiment each islet<br>preparation was used) | Islet dispersion and<br>flow cytometry | Islet dispersion and<br>flow cytometry | Islet dispersion and<br>flow cytometry |

**Supplemental Figure 1.**
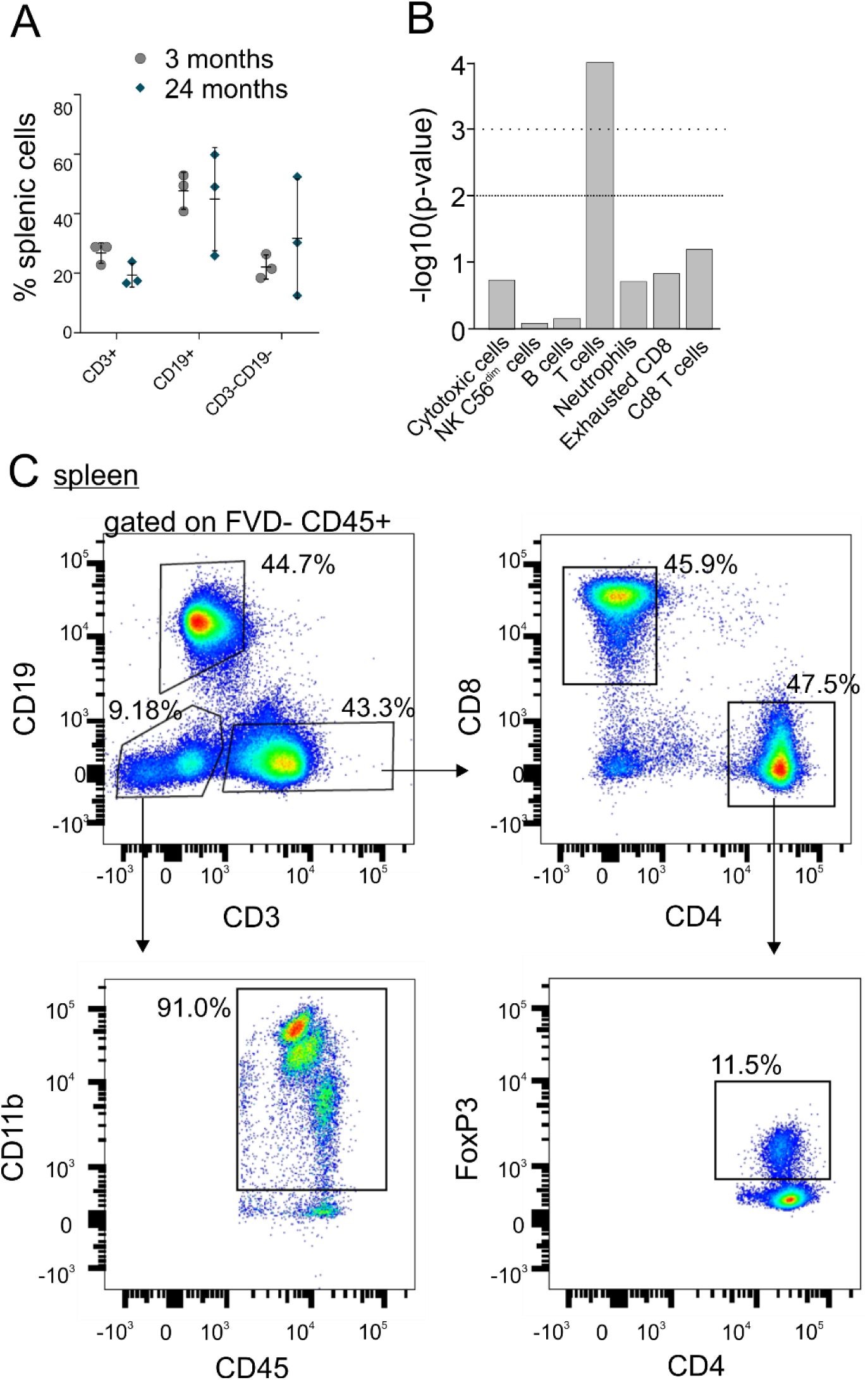
A) Quantification of immune cell populations as a percent of viable cells in spleens from 3- and 24-month old mice, n=3, gated on FSC, SSC, viability (7AAD-) and CD45+. B) Cell profile analysis was performed on transcriptome profiles obtained from sorted CD3+ islet cells from 3- and 24-month old mice via NSolver4.0, revealing a statistically significant T cell profile relative to other cell types, n=3 samples, 5-7 mice pooled per sample. C) Spleen from a 10-month old mouse was dispersed for analysis and gated on FSC, SSC, viability (FVD-) and CD45+, and subsequently CD19, CD3, CD11b, CD8, CD4 and FoxP3. Numbers in FACS plots represent the percent of cells in each selection as a function of the parent population.

**Supplemental Figure 2.**
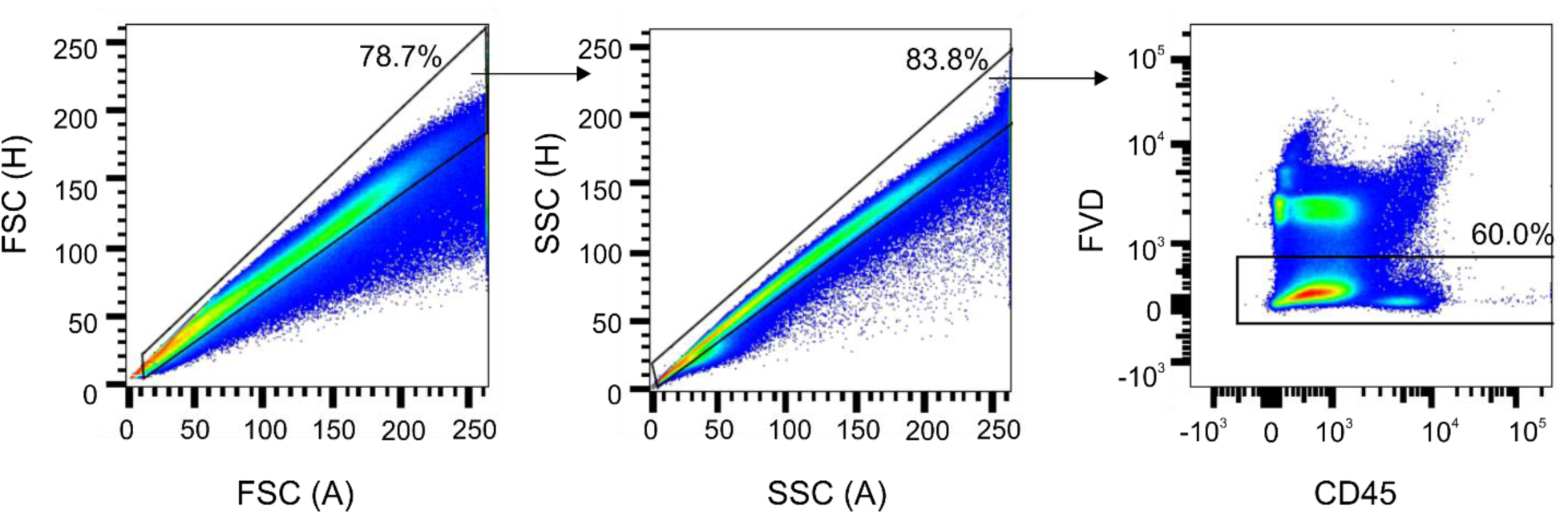
Representative data from human islet preparations, dispersed and gated on FSC, SSC and viability (FVD-). FACS plots are representative of 3 independent biological samples. Numbers in FACS plots represent the percent of cells in each selection as a function of the parent population.

